# Draft Genome of the Sea Cucumber Holothuria glaberrima, a model for the study of regeneration

**DOI:** 10.1101/2020.05.26.117028

**Authors:** Joshua G. Medina-Feliciano, Stacy Pirro, Jose E. García-Arrarás, Vladimir Mashanov, Joseph F. Ryan

**Affiliations:** Biology Department, University of Puerto Rico, Río Piedras, Puerto Rico, 00931, USA; Iridian Genomes, Inc., Bethesda, MD, 20817, USA; Wake Forest Institute for Regenerative Medicine, Winston Salem, NC, USA; Whitney Laboratory for Marine Bioscience, University of Florida, St. Augustine, FL, 32080, USA

**Keywords:** holothuroid, echinoderm, regeneration, melanotransferrin, genome assembly, mitochondrial genome

## Abstract

Regeneration is one of the most fascinating and yet least understood processes of animals. Echinoderms, one of the closest related invertebrate groups to humans, can contribute to our understanding of the genetic basis of regenerative processes. Amongst echinoderms, sea cucumbers have the ability to grow back most of their body parts following injury, including the intestine and nervous tissue. The cellular and molecular events underlying these abilities in sea cucumbers have been most extensively studied in the species *Holothuria glaberrima*. However, research into the regenerative abilities of this species have been impeded due to the lack of adequate genomic resources. Here, we report the first draft genome assembly of *H. glaberrima* and demonstrate its value for future genetic studies. Using only short sequencing reads, we assembled the genome into 2,960,762 scaffolds totaling 1.5 gigabases with an N50 of 15 kilobases. Our BUSCO assessment of the genome resulted in 882 (90.2%) complete and partial genes from 978 genes queried. We incorporated transcriptomic data from several different life history stages to annotate 41,076 genes in our final assembly. To demonstrate the usefulness of the genome, we fully annotated the melanotransferrin (*Mtf)* gene family, which have a potential role in regeneration of the sea cucumber intestine. Using these same data, we extracted the mitochondrial genome, which showed high conservation to that of other holothuroids. Thus, these data will be a critical resource for ongoing studies of regeneration and other studies in sea cucumbers.

## Introduction

Regeneration, the replacement of lost or damaged body parts, is one of the most fascinating processes of animals. Regenerative abilities are widespread in the animal kingdom [1], but our understanding of the evolution of regenerative capacity is based primarily on research into a small number of animal species. Echinoderms, along with hemichordates make up the clade Ambulacraria, the sister lineage to chordates [2]. Unlike chordates, many echinoderms, especially sea cucumbers, have extensive regenerative abilities. Nevertheless, relatively few genomic resources exist for sea cucumber models [3, 4].

Sea cucumbers (holothurians) have the ability to regenerate their intestine following evisceration [5] and their radial nerve cord following transection [6]. Many cellular events that underlie these abilities have been best described in the species *Holothuria glaberrima*. For instance, details of cell migration, dedifferentiation, division and apoptosis during regeneration of the digestive tract have all been published [7]. Dedifferentiation, for example, is known to occur after evisceration in the mesentery muscle layer that is adjacent to the injury site [8, 9]. While we know exactly where and when these events occur at the cellular level, the underlying molecular infrastructure has not yet been clearly elucidated [10]. In recent years, we have reported gene expression profiles of various stages of regeneration, providing information on which genes play important roles in sea cucumber regenerative processes [11, 12]. These RNA-Seq analyses have led to interesting findings, but the lack of a reference genome has limited the impact of these results.

Here we report the nuclear and mitochondrial genomes of *H. glaberrima*. We show the utility of our draft nuclear genome by using it to ascertain the genomic structure of melanotransferrins, which might play an important role in the holothurian regeneration process [13,14]. Melanotransferrins are membrane-bound molecules involved in many vertebrate processes and diseases, including development, cancer, and Alzheimer’s disease [15]. Moreover, in the sea cucumber, melanotransferrin genes are highly expressed in the intestine after immune activation [16]. We previously reported, based on transcriptomic data, that *H. glaberrima* contains four different melanotransferrin genes (HgMTF1, HgMTF2, HgMTF3, and HgMTF4), while there has been only one melanotransferrin reported in all other non-holothuroid animals sampled to date [14]. Furthermore, it has been hypothesized that the reason genomes from most species contain only one melanotransferrin is that multiple copies may act as a dominant lethal, preventing organisms with multiple copies from surviving [17]. The four melanotransferrin genes found in the *H. glaberrima* are highly expressed at different stages of intestinal regeneration. For these reasons, analyzing the genomic characteristics of these four melanotransferrin genes will shed light on the evolution of this gene family and will facilitate further studies on their specific role and regulation.

Comparisons of mitochondrial genomes, the remnants of an ancient endosymbiotic event that led to the origin of eukaryotes [18], provide insight into the evolution of aerobic respiration across the tree of life. When sequencing nuclear genomes, many copies of the mitochondrial genome are also sequenced. We therefore have assembled, annotated, and compared the mitochondrial genome of *H. glaberrima* with two other published holothuroid mitochondrial genomes.

The nuclear and mitochondrial genomes of *H. glaberrima* will be valuable resources for ongoing investigations into regenerative processes in this animal. In addition, these data will be useful for future studies in systematics, immunology, and evolution, among others.

## Methods and Results

### Data Availability, Reproducibility, and Transparency Statement

The raw genomic reads are available at NCBI’s Short Read Archive (Accession: SRR9695030). The genome assembly is available at http://ryanlab.whitney.ufl.edu/genomes/Holothuria_glaberrima/. Custom scripts, command lines, and data used in these analyses and alignment and tree files are available at https://github.com/josephryan/Holothuria_glaberrima_draft_genome. To maximize transparency and minimize confirmation bias, phylogenetic analyses were planned *a priori* in a phylotocol [19] which was posted to our GitHub repository (URL above). For these analyses we used the bridges system [20] part of the Extreme Science and Engineering Discovery Environment (XSEDE) [21].

### Data Collection

A single adult male sea cucumber (Figure 1) was collected from the northeast rocky coast of Puerto Rico (San Juan) and kept in aerated sea water until dissection. Gonads were collected into ethanol and sent to Iridian Genomes Inc. for DNA extraction, library preparation, and sequencing (Bethesda, MD). Of note, gonad tissue, which is a suboptimal source of DNA for genome sequencing due to potential crossover events, was used only after we made two failed attempts to use non-gonad tissues as a source of genomic DNA.

**Figure 1.**
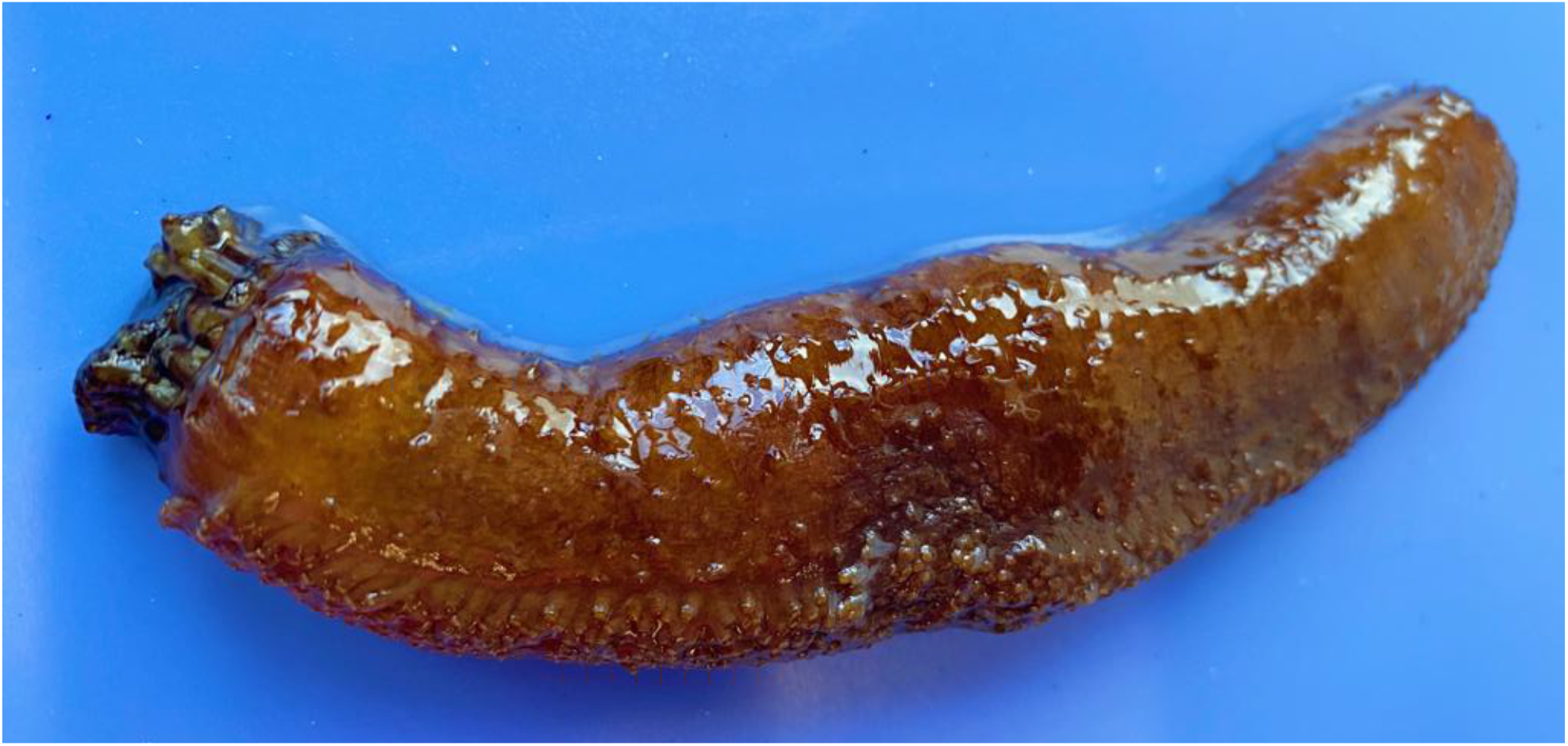
The sea cucumber *Holothuria glaberrima*.

### Pre-Assembly Processing

Sequencing was performed using an Illumina HiSeq X Ten instrument, which generated a total of 421,770,588 paired-end reads (126.5 gigabases) of length 150 nucleotides each. These reads were deposited in NCBI’s Short Read Archive (SRA) [22] under the accession SRR9695030 within one week of sequencing. Adapters from the raw sequencing reads were trimmed using Trimmomatic v0.39 (ILLUMINACLIP:2:30:10 LEADING:3 TRAILING:3 SLIDING-WINDOW:4:15 MINLEN:36) [23]. After Trimmomatic, 400,075,279 (95%) pairs were retained and 4,200,427,854 (3.5%) of bases making up these retained pairs were trimmed. We used ALLPATHS-LG v.44837 ErrorCorrectReads.pl script [24] to run error correction on reads that had been trimmed for adapters, in the process we removed a total of 1,104,390 read pairs (0.1%), orphaned (removed 1 from a pair) an additional 12,545,131 read pairs and made 117,247,709 corrections to 95,509,388 (11.6%) reads (an average of 1.2 corrections per corrected read). The genome size estimation generated by ErrorCorrectReads.pl was 1.2 Gb (1,171,092,004 bases) with a coverage estimate of 84X. This number is close to the estimate obtained with Feulgen densitometry (~1.4 Gb).

We next created filtered mitochondrial reads from the resulting trimmed/error-corrected reads. To do this we generated a preliminary assembly using Platanus Genome Assembler v1.2.4 [25] with k=45 and then performed a BLASTN search against this assembly using the *Holothuria scabra* mitochondrial genome as a query. We identified a single contig that represented the entire mitochondrial genome. We then used FastqSifter v1.1.3 [26] to remove all trimmed/error-corrected reads that aligned to the *H. glaberrima* mitochondrial genome. FastqSifter removed 53,304 read pairs and 3,583 unpaired reads, leaving 386,920,975 read pairs and 33,176,161 unpaired reads, which were used to assemble the final nuclear genome.

### Genome Assemblies and Annotation

We used Platanus Genome Assembler v1.2.4 using a range of kmer values (31, 45, 59, 73, and 87), from which K=31 resulted in the assembly with the highest N50 (11,332). In order to incorporate contiguous regions brought together in suboptimal assemblies that were absent from our K31 optimal assembly, we used Matemaker v1.0.0 [27] to generate artificial mate-pairs from our suboptimal assemblies (i.e., K45, K59, K73 and K87) with insert sizes of 2K, 5K and 10K. We then used SSPACE v3.0 [28] to scaffold the optimal K31 contig assembly using the generated mate-pairs. The resulting assembly consisted of 2,960,762 scaffolds comprised of 1,468,331,553 bases (close to the *in silico* and densitometry estimations above) with an N50 of 15,069. The longest scaffold was 244,111 nucleotides. There were 2,718,713 scaffolds shorter than 200 nucleotides. Many of these, but definitely not all, are likely errors.

We assessed the final assembly with BUSCO v3 [29] against the metazoan database consisting of 978 genes using the web assessment tool gVolante [30]. Our BUSCO assessment resulted in recovering 731 (74.7%) complete and 151 (15.4%) partial genes from the 978 queried genes in the Metazoa dataset (Table 1).

**Table 1.**
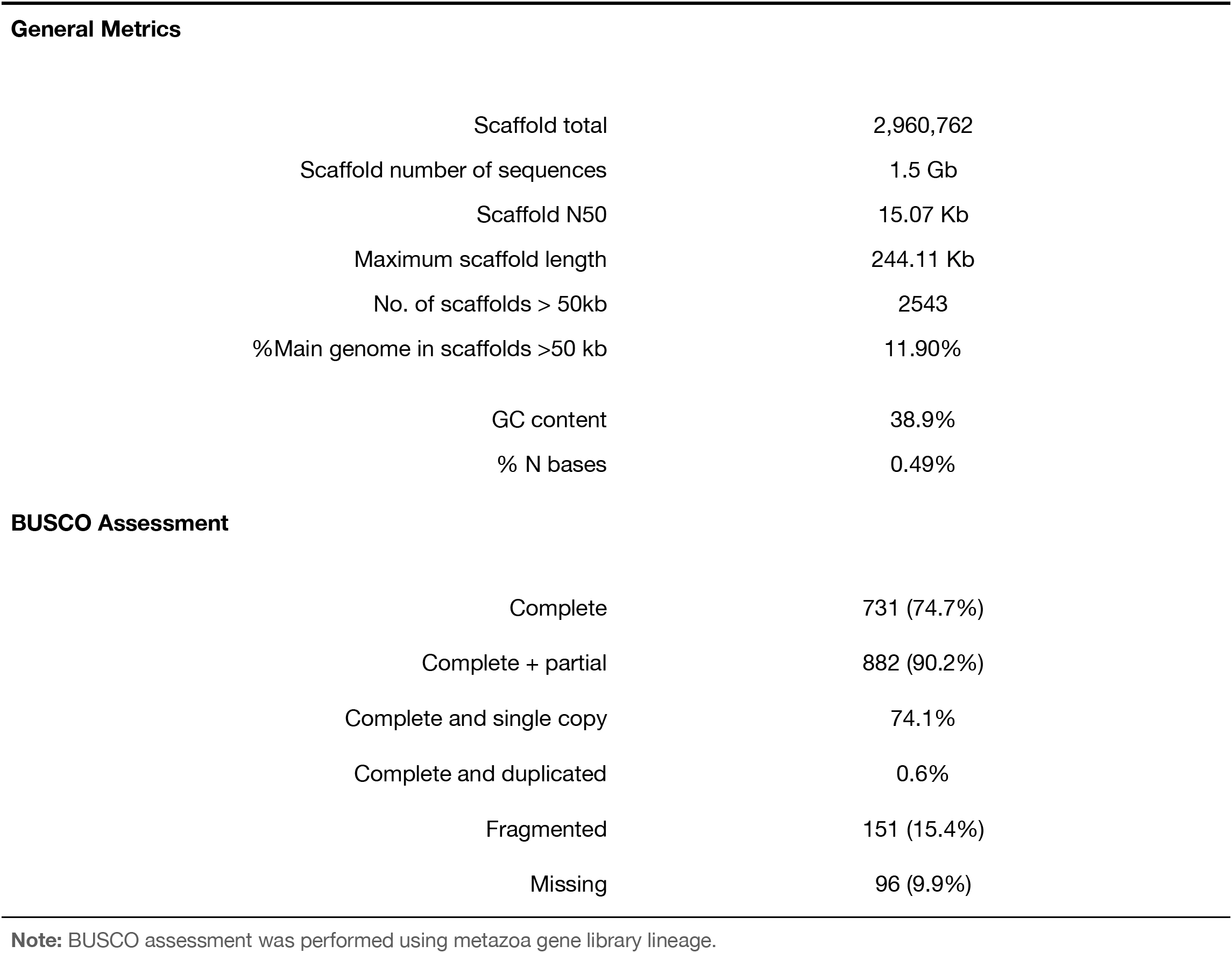
General Genome Scaffold Assembly Metrics and BUSCO Completeness Assessment.

|  |  |
| --- | --- |
| Scaffold total | 2,960,762 |
| Scaffold number of sequences | 1.5 Gb |
| Scaffold N50 | 15.07 Kb |
| Maximum scaffold length | 244.11 Kb |
| No. of scaffolds > 50kb | 2543 |
| %Main genome in scaffolds >50 kb | 11.90% |
| GC content | 38.9% |
| % N bases | 0.49% |
| **BUSCO Assessment** |  |
| Complete | 731 (74.7%) |
| Complete + partial | 882 (90.2%) |
| Complete and single copy | 74.1% |
| Complete and duplicated | 0.6% |
| Fragmented | 151 (15.4%) |
| Missing | 96 (9.9%) |

**Note:** BUSCO assessment was performed using metazoa gene library lineage.

We used Augustus v3.3.3 [31] to predict gene models. As part of this process, we generated a hints file from our transcriptomic data (unpublished) by aligning our assembled transcriptome to our genome assembly using BLAT v35×1 [32], filtering these alignments with the Augustus utility script filter PSL.pl and then sorting the alignments. We next applied the Augustus utility scripts aln2wig, wig2hints.pl, and blat2hints.pl to create the final hints file for Augustus. In the final prediction step, we set the species parameter to *Strongylocentrotus purpuratus*. We predicted 41,076 gene models with a mean exon length of 185 bp and mean intron length of 1916 bp as calculated using GenomeFeatures v3.10 Bioconductor package [33]. Our BUSCO assessment of translated versions of these gene models resulted in the recovery of a total of 617 (63.1%) complete and 211 (21.6%) partial genes from the 978 queried genes in the Metazoa dataset (Table 2).

**Table 2.** AUGUSTUS Gene Model Prediction General Statistics.

| **Metric** | **Value** |
| --- | --- |
| Genes | 41,076 |
| Exons | 171,977 |
| Introns | 148,101 |
| Mean exon length | 185.38 |
| Mean intron length | 1,915.76 |
| **BUSCO assessment** |  |
| Complete + partial | 828 (84.7%) |
| Complete and single-copy | 568 (58.1%) |
| Complete and duplicated | 49 (5.0%) |
| Fragmented | 211 (21.6%) |
| Missing | 150 (15.3%) |

**Table 3.** *Holothuria glaberrima* mitochondrial genome features.

| Gene Name | Strand | Start | Stop | Length | Codon |  | Anticodon | Inter Nucl |
| --- | --- | --- | --- | --- | --- | --- | --- | --- |
|  |  |  |  |  | Start | Stop |  |  |
| *cox1* | + | 1 | 1557 | 1557 | ATG | TAA |  | 11 |
| tRNA-R | + | 1568 | 1634 | 67 |  |  | TCG | 0 |
| *nad4l* | + | 1635 | 1931 | 297 | ATG | TAA |  | 0 |
| *cox2* | + | 1932 | 2619 | 688 | ATG | T-- |  | 0 |
| tRNA-K | + | 2620 | 2686 | 67 |  |  | CTT | 0 |
| *atp8* | + | 2687 | 2851 | 165 | ATG | TAA |  | -7 |
| *atp6* | + | 2845 | 3531 | 687 | ATG | TAA |  | 2 |
| *cox3* | + | 3534 | 4316 | 783 | ATG | TAA |  | -2 |
| tRNA-S<sup>(UCA)</sup> | - | 4315 | 4385 | 71 |  |  | TGA | 18 |
| *nad3* | + | 4404 | 4748 | 345 | ATG | TAA |  | 0 |
| *nad4* | + | 4752 | 6116 | 1365 | ATG | TAG |  | -7 |
| tRNA-H | + | 6110 | 6179 | 70 |  |  | GTG | 1 |
| tRNA-S<sup>(AGC)</sup> | + | 6181 | 6248 | 68 |  |  | GCT | 0 |
| *nad5* | + | 6249 | 8081 | 1833 | ATG | TAA |  | 17 |
| *nad6* | - | 8099 | 8587 | 489 | ATG | TAG |  | 8 |
| *cob* | + | 8596 | 9738 | 1143 | ATG | TAA |  | -1 |
| tRNA-F | + | 9738 | 9808 | 71 |  |  | GAA | 0 |
| *srRNA* | + | 9809 | 10637 | 829 |  |  |  | 1 |
| tRNA-E | + | 10639 | 10707 | 69 |  |  | TTC | 0 |
| tRNA-T | + | 10708 | 10777 | 70 |  |  | TGT | 0 |
| PC-region | + | 10778 | 11049 | 272 |  |  |  | 0 |
| tRNA-P | + | 11050 | 11116 | 67 |  |  | TGG | -4 |
| tRNA-Q | - | 11113 | 11182 | 70 |  |  | TTG | 2 |
| tRNA-N | + | 11185 | 11254 | 70 |  |  | GTT | 0 |
| tRNA-L<sup>(CUA)</sup> | + | 11255 | 11326 | 72 |  |  | TAG | 5 |
| tRNA-A | - | 11332 | 11398 | 67 |  |  | TGC | 0 |
| tRNA-W | + | 11399 | 11466 | 68 |  |  | TCA | 0 |
| tRNA-C | + | 11467 | 11530 | 64 |  |  | GCA | 5 |
| tRNA-V | - | 11536 | 11605 | 70 |  |  | TAC | 20 |
| tRNA-M | + | 11626 | 11694 | 69 |  |  | CAT | 2 |
| tRNA-D | - | 11697 | 11766 | 70 |  |  | GTC | 1 |
| tRNA-Y | + | 11768 | 11835 | 68 |  |  | GTA | 0 |
| tRNA-G | + | 11836 | 11902 | 67 |  |  | TCC | 6 |
| tRNA-L<sup>(UUA)</sup> | + | 11909 | 11979 | 71 |  |  | TAA | 0 |
| *nad1* | + | 11980 | 12951 | 972 | ATG | TAA |  | 14 |
| tRNA-I | + | 12966 | 13033 | 68 |  |  | GAT | 0 |
| *nad2* | + | 13034 | 14077 | 1044 | ATG | TAA |  | 0 |
| *lrRNA* | + | 14078 | 15617 | 1540 |  |  |  | 0 |

### Comparative Genomics

We used OrthoFinder v2.2.7 [34] with default parameters to generate orthogroups using protein models from the newly assembled *H. glaberrima* genome, as well as from *Homo sapiens, Mus musculus, Danio rerio, and S. purpuratus* along with our *H. glaberrima* transcriptome that had been translated with TransDecoder v. 5.3.0 [35] (NOTE: we included the translated *H. glaberrima* transcriptome in this analysis, because we realized that the *H. glaberrima* gene models will be incomplete due to the fragmented nature of the genome assembly; this should not be a problem for the other included datasets). The datasets for this analysis were obtained from the Ensembl FTP database (*H. sapiens, M. musculus, D. rerio*) and from the EchinoBase (*S. purpuratus*). Our OrthoFinder analysis resulted in a total of 52,812 orthogroups. We identified 5,918 orthogroups shared between all species, 8,974 that contained only sequences from *H. glaberrima* and the purple sea urchin *S. purpuratus*, and close to 7,000 for each of the groups that included only *H. glaberrima* and one of the vertebrates (Figure 2A).

**Figure 2.**
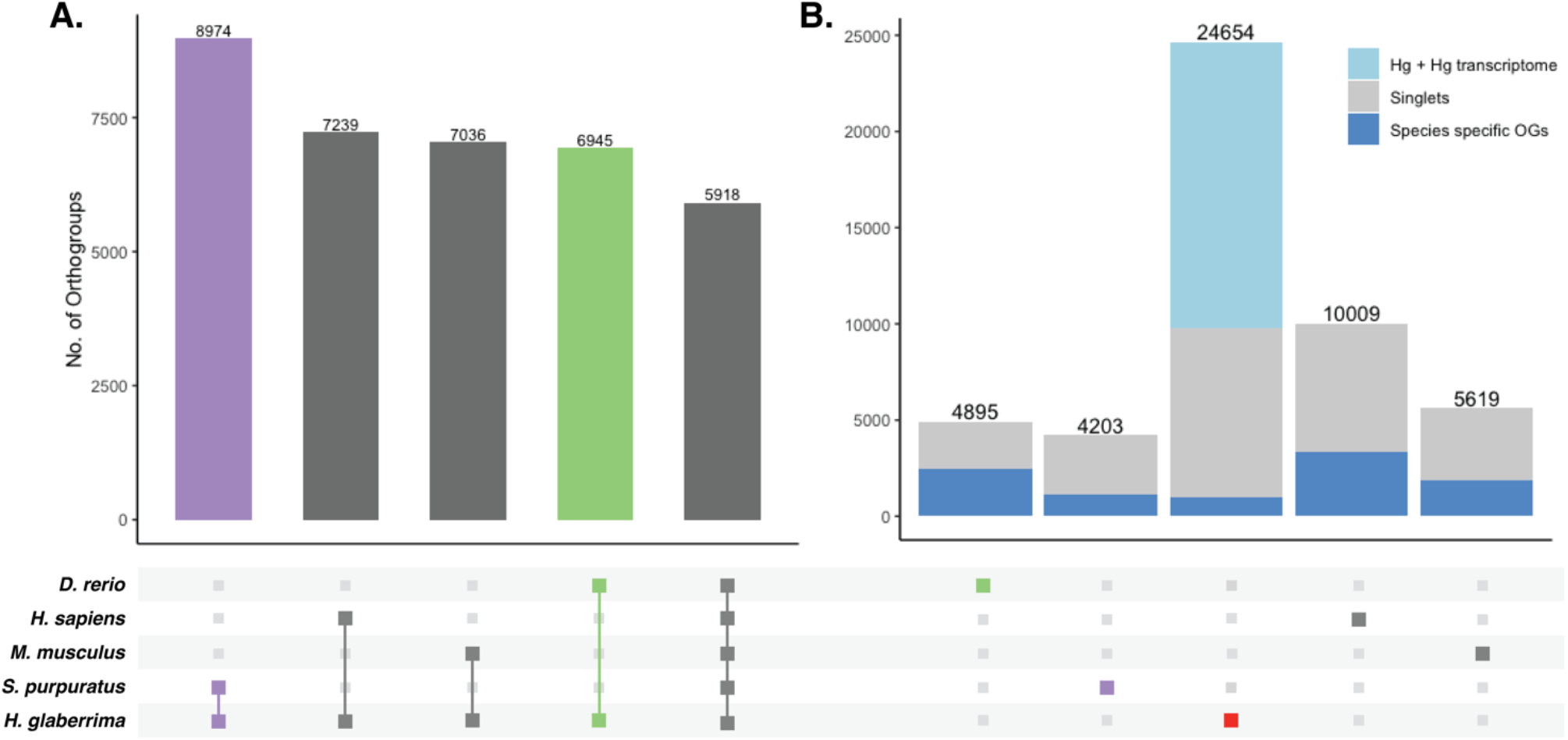
Results of Orthogroups from OrthoFinder Analysis. OrthoFinder analyses were performed with amino acid sequences of *H. glaberrima* protein models (based on predicted gene models), *H. glaberrima* transcriptome, *D. rerio, H. sapiens, M. musculus*, and *S. Purpuratus*. (A) Total orthogroup overlap between *H. glaberrima* and each other species as well as total orthogroups shared between all species. (B) Species-specific orthogroups for all species analyzed. These include singlets –sequences that didn’t form a group with any other sequence in the analysis and orthogroups consisting of only sequences from a single species. Since we incorporate both translated transcripts and protein models for *H. glaberrima*, orthogroups that include these two categories (Hg + Hg transcriptome) but no other species are included in our tally of *H. glaberrima* species-specific orthogroups. Bottom panel represents pattern of corresponding species in each bar of top plot.

We identified 24,654 *H. glaberrima*-specific orthogroups; this included those that consisted solely of *H. glaberrima* protein models (i.e. protein sequences derived from translated genomic sequences), those that included both *H. glaberrima* protein models and translated transcripts, and those consisting of only a single *H. glaberrima* protein model (“singlets”). We did not count those that included only *H. glaberrima* translated transcripts as many of these were spurious translations (by design to maximize sensitivity). There were far more species-specific orthogroups in *H. glaberrima* than there were in any of the other species—*H. sapiens* was next with 10,009 (Figure 2B).

### Melanotransferrin

We further evaluated our genome assembly by closely examining *H. glaberrima* genes that had been previously described in studies based on expressed sequence tags (EST), RNA sequencing, and/or Sanger-based cDNA sequencing. This approach allowed us to manually assess the accuracy of our genome assembly and to determine gene structures. Specifically, we selected the melanotransferrin gene family which is significantly upregulated in *H. glaberrima* intestinal tissue during intestine regeneration [13]. This group included *HgMtf1* (NCBI: GQ243222), *HgMtf2* (NCBI: KP861416), *HgMtf3* (NCBID: KP861417), *HgMtf4* (NCBI: KP861418).

We identified four *Mtf* genes in our genome assembly using NCBI BLASTN [36] (Figure 3A). The human Mtf gene (MELTF) is large, coding for 749 amino acids and its 16 exons span more than 28 kilobases in the human genome. Likewise, the *H. glaberrima* Mtf genes are similarly large (HgMtf1=739 aa, HgMtf2=723 aa, HgMtf3=752 aa, HgMtf4=750 aa), with all four genomic loci having 14 exons and spanning many kilobases. Due to the large size of these genes and the fragmented nature of our assembly, all four of the Mtf genes occur on multiple scaffolds (Figure 3A). Nevertheless, we were able to account for all exons and capture the structure of each gene. Besides differing from the human Mtf gene in the number of exons, there is variation in the position of several introns suggesting that there have been substantial structural changes in these genes since the last common ancestor of vertebrates and echinoderms.

**Figure 3.**
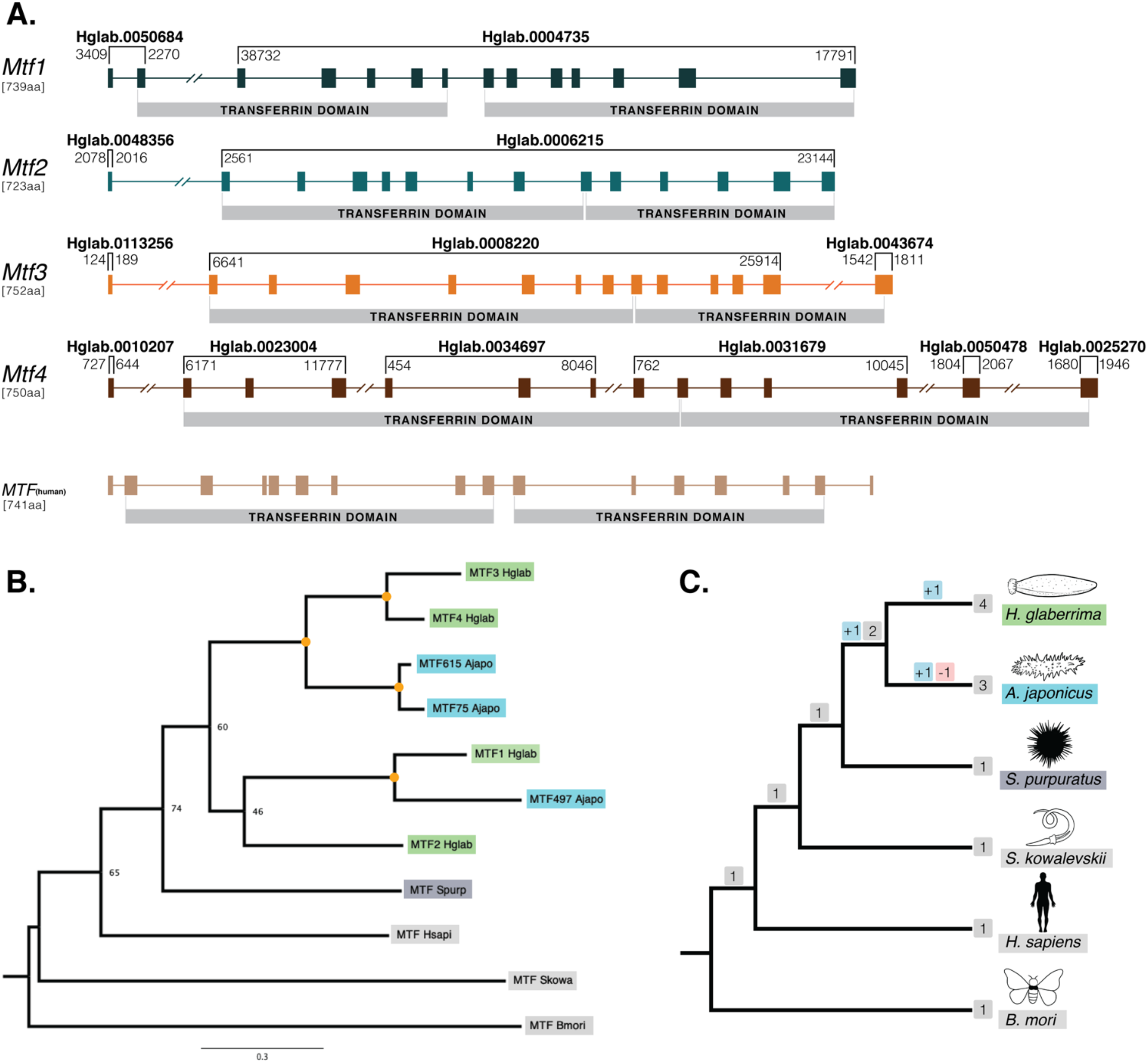
Gene structure and Phylogenetic Analysis of *H. glaberrima* melanotransferrin *(Mtf)* genes. (A) Genomic structure was determined using *H. glaberrima Mtf* gene and protein sequences present in NCBI from previous studies as reference [12]. Scaffolds containing specific exons of the genes are shown at the top of each bracket (Hglab.XXXXX). Coordinates of the sequence in each scaffold are shown among brackets based on genome nucleotide sequence. Human Mtf gene structure was acquired from NCBI genome browser (MELTF NCBI: NM_005929.6). Transferrin domains are shown at the bottom of each gene. (B) Gene tree of all Mtf genes from *H. glaberrima*, *A. japonicus*, *S. purpuratus, H. sapiens, S. kowalevskii*, and *B. mori*. Mtf gene sequences of the last three species were acquired from NCBI. *A. japonicus* Mtf gene names represent the scaffold number where they are found in its genome assembly. Orange circles in nodes indicate bootstrap values of 100. (C) Species tree of organisms used in the phylogenetic analysis of the Mtf genes. Gene duplication and loss are shown in red and blue squares, respectively.

Our assembly captures considerable variability between the HgMtf1 and the three other *H. glaberrima* Mtf genes. For example, the sequence that codes for the N-terminal transferrin domains of HgMtf1 is interrupted by 5 introns, whereas the corresponding coding regions in HgMtf2–4 include 7 introns. Likewise, the sequence that codes for the C-terminal transferrin domain of HgMtf1 includes 6 introns, whereas these same coding regions in HgMtf2–4 includes 5 introns (Figure 3A). These results show that there have been considerable evolutionary changes in the genomic architecture HgMtf1, suggesting that HgMtf1 did not arise very recently.

The 5’ sequence of HgMtf1 in the genome differs from the previously reported cDNA sequence in GenBank (ACS74869). There appears to be a single nucleotide present in the genome that is absent from the previously reported sequence. This frameshift leads to 71 fewer amino acids in the translation of the genomic HgMtf1. We surmise that the genome is correct and that the previously published sequence includes an erroneous single-nucleotide deletion based on the following evidence. A BLASTP of the 71 amino-acid portion against the GenBank Non-redundant database produces only a single match to itself. The region in question is exactly the same in all of our assemblies, including suboptimal assemblies. Moreover, TBLASTN of the 71 amino acids against our transcriptome produced no hits.

To deduce the relationship between the four paralogous *H. glaberrima* Mtf genes with those of other animals, we conducted a phylogenetic analysis of Mtf genes (Figure 3B-C). We used Mtf genes from *H. glaberrima*, *Apostichopus japonicus, S. purpuratus, Saccoglossus kowalevskii, H. sapiens*, and *Bombyx mori*. Only *A. japonicus*, the other sea cucumber in our analysis, included more than one Mtf gene in its genome [4]. Initial pre-genomic studies reported only one melanotransferrin in *A. japonicus* [37], but the latest genome data of this sea cucumber showed that there were multiple copies in its genome. We searched the predicted gene models of *A. japonicus* and found three Mtf genes, which we included in our phylogenetic analysis. Our criteria for inclusion was the presence of two full transferrin domains. It is possible that there are other Mtf genes present in the *A. japonicus* genome that were either missed or mispredicted in the current assembly. There is no other evidence to date showing more than one *Mtf* gene in the genomes of non-holothuroid species. As such, our phylogenetic analysis supports the hypothesis that *Mtf* gene family expansion is specific to the holothuroid lineage.

### Mitochondrial Genome

While assembling the *H. glaberrima* nuclear genome, we also assembled the mitochondrial genome. There was a 32-nucleotide overlap between the beginning and the end of the mitochondrial scaffold as would be expected from a circular genome. As such, we removed the overlap and reordered the scaffold so that the beginning of the scaffold corresponded with the beginning of the *cox1* gene. We used Mitos [38] to predict genes and tRNAscan-SE 2.0 [39] to annotate tRNAs. Every annotated gene was additionally confirmed with BLAST against *Holothuria scabra* mitochondrial genes. This approach resulted in more precise annotation as some of the stop codons were wrongly predicted by Mitos. Furthermore, tRNAscan-SE failed to predict some of the tRNAs in the *H. glaberrima* mitochondrial genome, in which case we used Mitos predictions.

The *H. glaberrima* mitochondrial genome is 15,617 bp, a similar size to that of previously reported holothuroids (*H. scabra*=15,779 bp, *H. leucospilata*=15,904 bp) [40]. Likewise, mitochondrial genome features of *H. glaberrima* showed the same gene order reported for *H. scabra* and similar characteristics to those of *H. scabra* and *H. leucospilata* [41,42]. Differences included the length of the putative control region (PC region) which was significantly smaller in *H. glaberrima* with 272 base pairs compared to 456 bp and 551 bp for *H. scabra* and *H. leucospilata*, respectively. Moreover, the total number of intergenic nucleotides was similar in *H. glaberrima* with 113 bp as *H. scabra* which contains 109 bp. Total intergenic nucleotides were distributed among 15 intergenic spacers, ranging from 1 to 20 bp in size, comparable to *H. scabra* and *H. leucospilata* which contain 15 and 16 intergenic spacers, respectively. There were 5 regions with overlapping genes within the genome, the most significant being those between *atp8* and *atp6*, and *nad4* and tRNA-His, with overlaps of 7 bp, followed by tRNA-P and tRNA-Q with overlap of 4 bp. From the 5 overlapping regions, *H. glaberrima* shares all the mentioned overlapping areas with both *H. scabra* and *H. leucospilata* suggesting that these are ancient overlaps. In addition, as reported for both *H. scabra* and *H. forskali, H. glaberrima* also has an incomplete stop codon T of *cox2*, supporting the likeliness of it being a distinctive feature of Holothuria [40,41].

## Conclusions

We sequenced, assembled, and annotated the first draft genome assembly of *H. glaberrima*, one of the best studied regenerative invertebrates at the morphological and cellular levels. We demonstrated the utility of the genome by analyzing the Mtf gene family genomic loci. We also assembled and annotated the *H. glaberrima* mitochondrial genome showing conservation with other mitochondrial genomes from the genus *Holothuria*. Our work provides a key resource for the sea cucumber *H. glaberrima*, which will be important for future studies in animal regeneration and beyond.

## Acknowledgements

The project was partially funded by the National Institute of Health (NIH) (Grant R15NS01686 and R21AG057974) and the University of Puerto Rico. Also, funding support was provided by National Science Foundation (NSF) [1542597 to J.F.R]; [1826558 to J.G.M.F]. This work used the Extreme Science and Engineering Discovery Environment (XSEDE), which is supported by NSF grant number ACI-1548562. Specifically, it used the Bridges system, which is supported by NSF award number ACI-1445606, at the Pittsburgh Supercomputing Center (PSC). Additionally, it used the High Performance Computing Facility of the University of Puerto Rico, sponsored by the University of Puerto Rico and the Institutional Development Award (IDeA) INBRE grant P20 GM103475 from the National Institute for General Medical Sciences (NIGMS), a component of the National Institutes of Health (NIH) and the Bioinformatics Research Core of INBRE.

## References

1. Bely AE, Nyberg KG. Evolution of animal regeneration: re-emergence of a field. Trends in Ecology & Evolution. 2010;25:161–70.

2. Cannon JT, Vellutini BC, Smith J, Ronquist F, Jondelius U, Hejnol A. Xenacoelomorpha is the sister group to Nephrozoa. Nature. 2016;530:89–93.

3. Long KA, Nossa CW, Sewell MA, Putnam NH, Ryan JF. Low coverage sequencing of three echinoderm genomes: the brittle star Ophionereis fasciata, the sea star Patiriella regularis, and the sea cucumber Australostichopus mollis. GigaSci. 2016;5:20.

4. Zhang X, Sun L, Yuan J, Sun Y, Gao Y, Zhang L, et al. The sea cucumber genome provides insights into morphological evolution and visceral regeneration. Tyler-Smith C, editor. PLOS Biology. 2017;15:e2003790.

5. García-Arrarás JE, Estrada-Rodgers L, Santiago R, Torres II, Díaz-Miranda L, Torres-Avillán I. Cellular mechanisms of intestine regeneration in the sea cucumber, Holothuria glaberrima Selenka (Holothuroidea:Echinodermata). Journal of Experimental Zoology. 1998;281:288–304.

6. San Miguel-Ruiz JE, Maldonado-Soto AR, García-Arrarás JE. Regeneration of the radial nerve cord in the sea cucumber Holothuria glaberrima. BMC Dev Biol. 2009;9:3.

7. García-Arrarás JE, Lázaro-Peña MI, Díaz-Balzac CA. Holothurians as a Model System to Study Regeneration. In: Kloc M, Kubiak JZ, editors. Marine Organisms as Model Systems in Biology and Medicine [Internet]. Cham: Springer International Publishing; 2018 [cited 2019 Mar 13]. p. 255–83. Available from: http://link.springer.com/10.1007/978-3-319-92486-1_13

8. García-Arrarás JE, Valentín-Tirado G, Flores JE, Rosa RJ, Rivera-Cruz A, San Miguel-Ruiz JE, et al. Cell dedifferentiation and epithelial to mesenchymal transitions during intestinal regeneration in H. glaberrima. BMC Dev Biol. 2011;11:61.

9. García-Arrarás JE, Bello SA, Malavez S. The mesentery as the epicenter for intestinal regeneration. Seminars in Cell & Developmental Biology. 2019;92:45–54.

10. Mashanov VS, García-Arrarás JE. Gut Regeneration in Holothurians: A Snapshot of Recent Developments. The Biological Bulletin. 2011;221:93–109.

11. Ortiz-Pineda PA, Ramírez-Gómez F, Pérez-Ortiz J, González-Díaz S, Santiago-De Jesús F, Hernández-Pasos J, et al. Gene expression profiling of intestinal regeneration in the sea cucumber. BMC Genomics. 2009;10:262.

12. Mashanov VS, Zueva OR, García-Arrarás JE. Transcriptomic changes during regeneration of the central nervous system in an echinoderm. BMC Genomics. 2014;15:357.

13. Rojas-Cartagena C, Ortíz-Pineda P, Ramírez-Gómez F, Suárez-Castillo EC, Matos-Cruz V, Rodríguez C, et al. Distinct profiles of expressed sequence tags during intestinal regeneration in the sea cucumber Holothuria glaberrima. Physiological Genomics. 2007;31:203–15.

14. Hernández-Pasos J, Valentín-Tirado G, García-Arrarás JE. Melanotransferrin: New Homolog Genes and Their Differential Expression during Intestinal Regeneration in the Sea Cucumber Holothuria glaberrima: MELANOTRANSFERRIN HOMOLOGS IN HOLOTHURIANS. J Exp Zool (Mol Dev Evol). 2017;328:259–74.

15. Rahmanto YS, Bal S, Loh KH, Yu Y, Richardson DR. Melanotransferrin: Search for a function. Biochimica et Biophysica Acta (BBA) - General Subjects. 2012;1820:237–43.

16. Ramírez-Gómez F, Ortiz-Pineda PA, Rivera-Cardona G, García-Arrarás JE. LPS-Induced Genes in Intestinal Tissue of the Sea Cucumber Holothuria glaberrima. Fugmann SD, editor. PLoS ONE. 2009;4:e6178.

17. Lambert LA, Perri H, Meehan TJ. Evolution of duplications in the transferrin family of proteins. Comparative Biochemistry and Physiology Part B: Biochemistry and Molecular Biology. 2005;140:11–25.

18. Martin WF, Garg S, Zimorski V. Endosymbiotic theories for eukaryote origin. Phil Trans R Soc B. 2015;370:20140330.

19. DeBiasse MB, Ryan JF. Phylotocol: Promoting Transparency and Overcoming Bias in Phylogenetics. Holder M, editor. Systematic Biology. 2019;68:672–8.

20. Nystrom NA, Levine MJ, Roskies RZ, Scott JR. Bridges: A uniquely flexible HPC resource for new communities and data analytics. Proceedings of the 2015 XSEDE conference: Scientific advancements enabled by enhanced cyberinfrastructure. New York, NY, USA: ACM; 2015. p. 30:1–30:8. Available from: http://doi.acm.org/10.1145/2792745.2792775

21. Towns J, Cockerill T, Dahan M, Foster I, Gaither K, Grimshaw A, et al. XSEDE: Accelerating scientific discovery. Computing in Science & Engineering. 2014;16:62–74.

22. Leinonen R, Sugawara H, Shumway M, on behalf of the International Nucleotide Sequence Database Collaboration. The Sequence Read Archive. Nucleic Acids Research. 2011;39:D19–21.

23. Bolger AM, Lohse M, Usadel B. Trimmomatic: a flexible trimmer for Illumina sequence data. Bioinformatics. 2014;30:2114–20.

24. Gnerre S, MacCallum I, Przybylski D, Ribeiro FJ, Burton JN, Walker BJ, et al. High-quality draft assemblies of mammalian genomes from massively parallel sequence data. Proceedings of the National Academy of Sciences. 2011;108:1513–8.

25. Kajitani R, Toshimoto K, Noguchi H, Toyoda A, Ogura Y, Okuno M, et al. Efficient de novo assembly of highly heterozygous genomes from whole-genome shotgun short reads. Genome Research. 2014;24:1384–95.

26. Ryan J. FastqSifter [Internet]. 2015. Available from: https://github.com/josephryan/FastqSifter

27. Ryan J. matemaker [Internet]. 2015. Available from: https://github.com/josephryan/matemaker

28. Boetzer M, Henkel CV, Jansen HJ, Butler D, Pirovano W. Scaffolding pre-assembled contigs using SSPACE. Bioinformatics. 2011;27:578–9.

29. Simão FA, Waterhouse RM, Ioannidis P, Kriventseva EV, Zdobnov EM. BUSCO: assessing genome assembly and annotation completeness with single-copy orthologs. Bioinformatics. 2015;31:3210–2.

30. Nishimura O, Hara Y, Kuraku S. gVolante for standardizing completeness assessment of genome and transcriptome assemblies. Hancock J, editor. Bioinformatics. 2017;33:3635–7.

31. Stanke M, Waack S. Gene prediction with a hidden Markov model and a new intron submodel. Bioinformatics. 2003;19:ii215–25.

32. Kent WJ. BLAT—The BLAST-Like Alignment Tool. Cold Spring Harbor Laboratory Press. 2002;

33. Lawrence M, Huber W, Pagès H, Aboyoun P, Carlson M, Gentleman R, et al. Software for Computing and Annotating Genomic Ranges. Prlic A, editor. PLoS Comput Biol. 2013;9:e1003118.

34. Emms DM, Kelly S. OrthoFinder: phylogenetic orthology inference for comparative genomics. Genome Biol. 2019;20:238.

35. Haas B, Papanicolaou A. Transdecoder [Internet]. 2018. Available from: https://github.com/TransDecoder/TransDecoder

36. AltschuP SF, Gish W, Miller W, Myers EW, Lipman DJ. Basic Local Alignment Search Tool. :8.

37. Qiu X, Li D, Cui J, Liu Y, Wang X. Molecular cloning, characterization and expression analysis of Melanotransferrin from the sea cucumber Apostichopus japonicus. Mol Biol Rep. 2014;41:3781–91.

38. Bernt M, Donath A, Jühling F, Externbrink F, Florentz C, Fritzsch G, et al. MITOS: Improved de novo metazoan mitochondrial genome annotation. Molecular Phylogenetics and Evolution. 2013;69:313–9.

39. Chan PP, Lowe TM. tRNAscan-SE: Searching for tRNA Genes in Genomic Sequences. In: Kollmar M, editor. Gene Prediction [Internet]. New York, NY: Springer New York; 2019 [cited 2020 Apr 12]. p. 1–14. Available from: http://link.springer.com/10.1007/978-1-4939-9173-0_1

40. Perseke M, Bernhard D, Fritzsch G, Brümmer F, Stadler PF, Schlegel M. Mitochondrial genome evolution in Ophiuroidea, Echinoidea, and Holothuroidea: Insights in phylogenetic relationships of Echinodermata. Molecular Phylogenetics and Evolution. 2010;56:201–11.

41. Xia J, Ren C, Yu Z, Wu X, Qian J, Hu C. Complete mitochondrial genome of the sandfish Holothuria scabra (Holothuroidea, Holothuriidae). Mitochondrial DNA Part A. 2016;27:4174–5.

42. Yang Q, Lin Q, Wu J, Tran NT, Huang R, Sun Z, et al. Complete mitochondrial genome of *Holothuria leucospilata* (Holothuroidea, Holothuriidae) and phylogenetic analysis. Mitochondrial DNA Part B. 2019;4:2751–2.

